# A Universal Chemical Method for Rational Design of Protein-based Nanoreactors

**DOI:** 10.1101/2021.03.01.433350

**Authors:** Mullapudi Mohan Reddy, Punita Bathla, Britto S. Sandanaraj

**Affiliations:** Department of Chemistry, Indian Institute of Science Education and Research - Pune; Department of Biology, Indian Institute of Science Education and Research - Pune

## Abstract

Self-assembly of a monomeric protease to form a multi-subunit protein complex “proteasome” enables targeted protein degradation in living cells. The naturally occurring proteasomes serve as an inspiration and blueprint for the design of artificial protein-based nanoreactors. Here we disclose a general chemical strategy for the design of proteasome-like nanoreactors. Micelle-assisted protein labeling (MAPLab) technology along with the N-terminal bioconjugation strategy is utilized for the synthesis of a well-defined monodisperse self-assembling semi-synthetic protease. The designer protein is programmed to self-assemble into a proteasome-like nanostructure which preserves the functional properties of native protease.

---

The proteasome machinery which is present in all three domains of life is the finest example for self-assembled protein-based nanostructure that carries out the essential function such as targeted protein degradation.^1^ In bacterial proteasome, the protease activity is performed by the *N*-terminal threonine residue of each subunit. The β-subunit of 14-mer is arranged in such a way that the *N*-terminal active residue is positioned inside the central cavity of the hollow cylindrical 20-S core particle to perform proteolysis of target substrates.^2^ This highly complex and most sophisticated nanoreactor is the result of billion years of evolution and therefore it is highly challenging to design an artificial proteasome with such complexity. Nevertheless, these naturally occurring nanomachines serve as a blueprint for the design of primitive proteasomes.

Encapsulation of proteases inside protein-based nanocompartments results in compartmentalization.^3^ This strategy is widely used in the living system for controlling substrate specificity and preventing unwanted off-target reactions.^4^ In an elegant study, Hilvert and coworkers demonstrated the encapsulation of a protease inside cage-forming lumazine synthase and achieved artificial substrate selectivity.^5^ This top-down approach is also exploited by other groups for the design of various protein-based nanoreactors.^6^ Although extremely powerful, these methods rely on the engineering of naturally occurring protein cages through directed evolution and are therefore restricted to a few scaffolds. In contrast to the top-down strategy, the bottom-up approaches provide opportunities to design protein assemblies with an extraordinary diversity. In this aspect, computational protein design has proven to be a mature technology over the past decade.^7^ However, most of the reports are based on the design of static protein assemblies,^8^ and to our knowledge there are very few reports on synthetic protein nanoassemblies that can exhibit protease activity.^9^

Chemical technologies are emerging as a complementary method for the design of protein assemblies.^10^ Recently, our group invented a new method named “micelle-assisted protein labeling (MAPLab) technology (Figure 1a).^11^ This technology was used for the design of a diverse set of self-assembling semi-synthetic proteins which includes protein amphiphiles^11a^, protein-dendron bioconjugates^11b^, protein-peptide bioconjugates^11c^, photo-responsive protein amphiphiles,^11a^ and photo-sensitive protein-dendron bioconjugates^11d^. We have demonstrated the successful design of monodisperse protein nanoparticles with a controlled oligomeric state, tunable size, and surface charges.^11a^ We also demonstrated the design of a multi-responsive protein nanoparticle that can disassemble upon exposure to different stimuli.^11a, d^ However, in our previous method, the increased nucleophilicity of the active site residue of protein was exploited for site-specific bioconjugation and therefore the resultant protein assembly is catalytically inactive. Besides, the reported method is only applicable to the serine protease family. Although there are hundreds of enzymes that belong to this category^12^, however, the number of proteins is far less compared to the vast number of proteins that exist in Nature. Therefore, it is important to develop a general chemical methodology that would work for the vast majority of the proteins to make self-assembling semi-synthetic proteins without affecting the function of the target protein. However, to the best of our knowledge, there are no chemical methods available that fulfill all of the above criteria.

**Figure 1.**
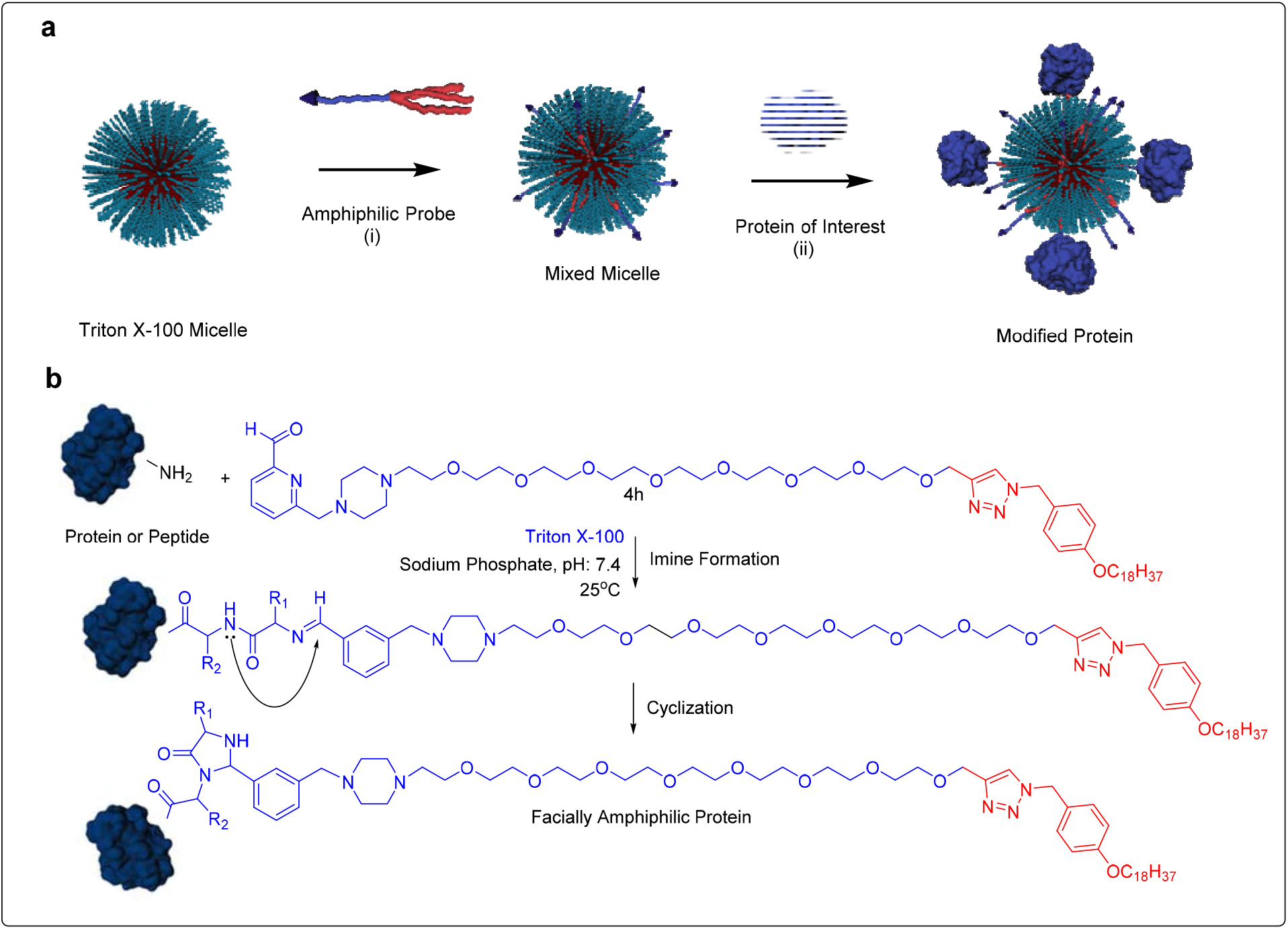
a. Schematic presentation of MAPLab technology. b. Protein bioconjugation reaction: custom-designed probe site-specifically reacts with *N*-terminal residue of the target protein to yield functional self-assembling semi-synthetic proteins.

During our research, we came across an exciting report from Francis and coworkers. They reported a new chemical methodology for site-specific modification of the *N*-terminal residue of a protein (Figure 1b).^13^ Unlike other bioconjugation methods, this method is versatile and can be used to label a large number of proteins irrespective of their size, shape, surface charge, molecular weight, and secondary, tertiary, and quaternary structures. Although a terrific method, one of the severe limitations of this methodology is that it is used only for bioconjugation of “hydrophilic payloads” which includes imaging agents, drugs, and polymers to make hydrophilic protein conjugates which find applications in the area of targeted drug delivery^14^ and biomaterials applications.^15^ Although extremely useful, this method cannot be directly used for site-specific bioconjugation of hydrophobic synthetic molecules or macromolecules.

Inspired by this work, we envisioned using this method along with MAPLab technology for the design of functional self-assembling semi-synthetic proteins (FSSSPs). Accordingly, we designed our target probe **8**. This probe contains three structural motifs: (i) 2-pyridine carboxyaldehyde as a site-specific *N*-terminal protein reactive group^13^ (ii) octaethylene linker as a hydrophilic linker, (iii) Octadeca benzyl group as a hydrophobic tail (Scheme 1). The choice of the linker and hydrophobic tail were based on our previous knowledge.^11a^ The synthesis of the target molecule was achieved in seven steps with an overall yield of 9.8%. In brief, intermediate **3** was synthesized following a procedure reported by Francis and coworkers.^13^ Similarly, intermediates **4** and **5** were made following synthetic procedures reported by our group.^11a^ Copper-catalyzed Huisgen [3+ 2] cycloaddition of compounds **4** and **5** yielded intermediate **6**.^11a^ The obtained intermediate **6** was reacted with piperazine in the presence of THF to get compound **7**. Finally, compound **7** upon treatment with compound **3** in the presence of K_2_CO_3_ in acetonitrile yielded the amphiphilic probe **8** (Scheme 1, see SI for details).

**Scheme 1:**
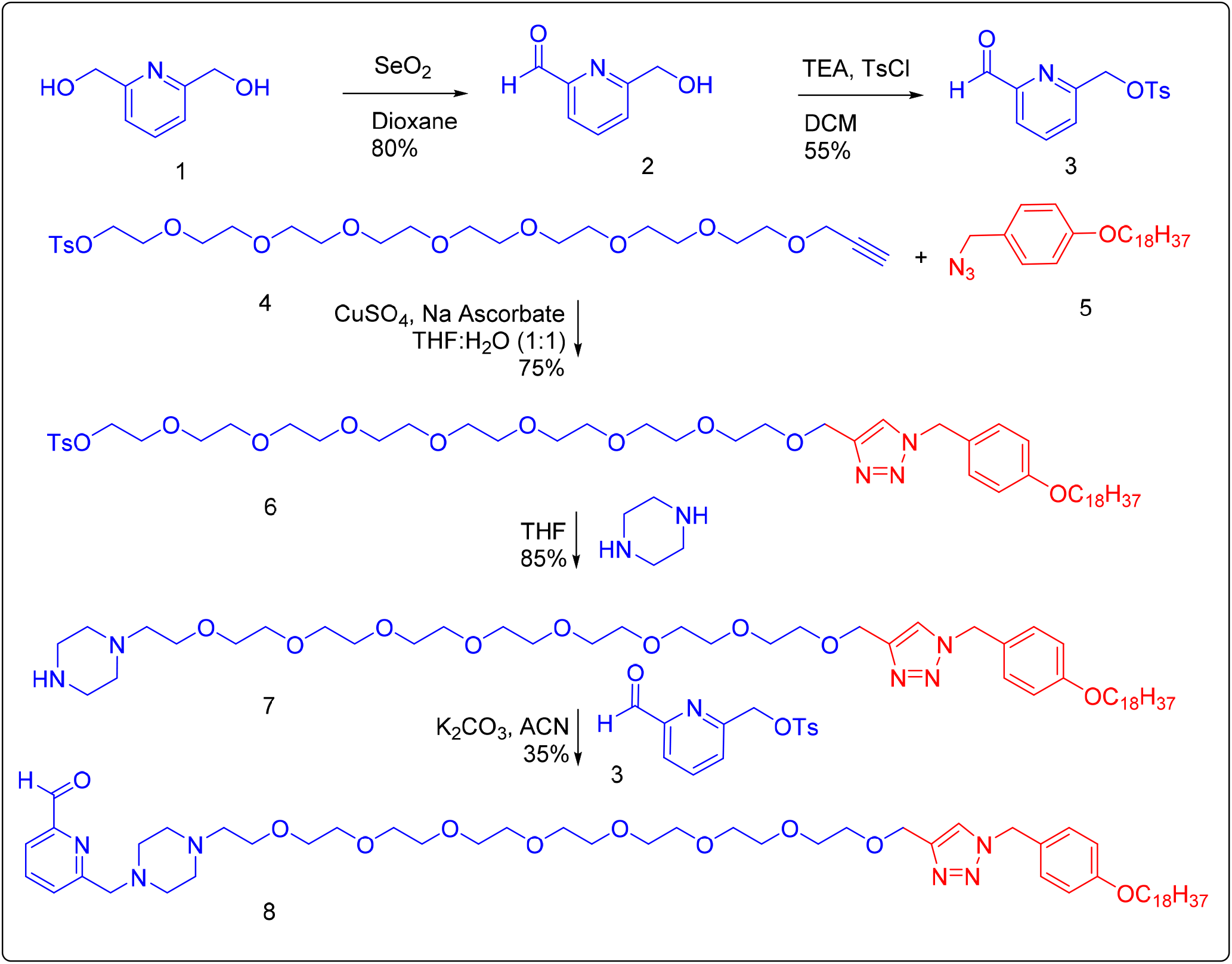
Multi-step synthesis of the target probe.

Considering the amphiphilic nature of the probe molecule, we tested the solubility of probe **8** in an aqueous medium. Unfortunately, probe **8** was not soluble presumably due to the presence of a long hydrophobic tail. Next, we tested whether the probe could be solubilized in the presence of neutral surfactant Triton-X-100. To our delight, the probe was soluble in the presence of Triton-X-100 because of mixed-micelle formation in which the protein-reactive group 2-pyridine carboxaldehyde is presumably presented outside the micellar surface.^11a-d^ Next, we carried out a bioconjugation reaction (Figure 1b) of probe **8** using a MAPLab technology pioneered by our group (Figure 1a).^11^ As a test case; we have attempted a bioconjugation reaction with a small globular protein lysozyme (abbreviated as LYS hereafter) in an aqueous medium at pH 7.2 room temperature for 12 h. The extent of bioconjugation reaction was monitored using a MALDI-ToF-MS. To our delight, we observed a new peak at high molecular weight corresponding to the facially amphiphilic bioconjugate LYS-OEG-C18 (Figure 2a). Note that this result indeed suggests that the 2-pyridine carboxaldehyde is present on the exterior surface of a mixed-micelle; otherwise, this reaction would not have been viable. Excited by these results, we wanted to investigate whether this reaction could be used for different categories of proteins. Accordingly, we chose three more proteins; a protease (chymotrypsin – abbreviated as CHY), a carrier protein (bovine serum albumin abbreviated as BSA), and a green fluorescent protein (GFP). All four proteins have different sizes, shapes, surface charges, secondary/tertiary structures and therefore modestly represent the structural and functional diversity of naturally occurring proteins.^13^ Bioconjugation of all three proteins was carried out similarly as described above and the progress of the reaction was monitored using the mass spec. To our delight, all of them are equally reactive as evident from mass spec results (Figure 2b-2d). As expected, we did not observe any double-labeling which is consistent with the published report.^13^

**Figure 2:**
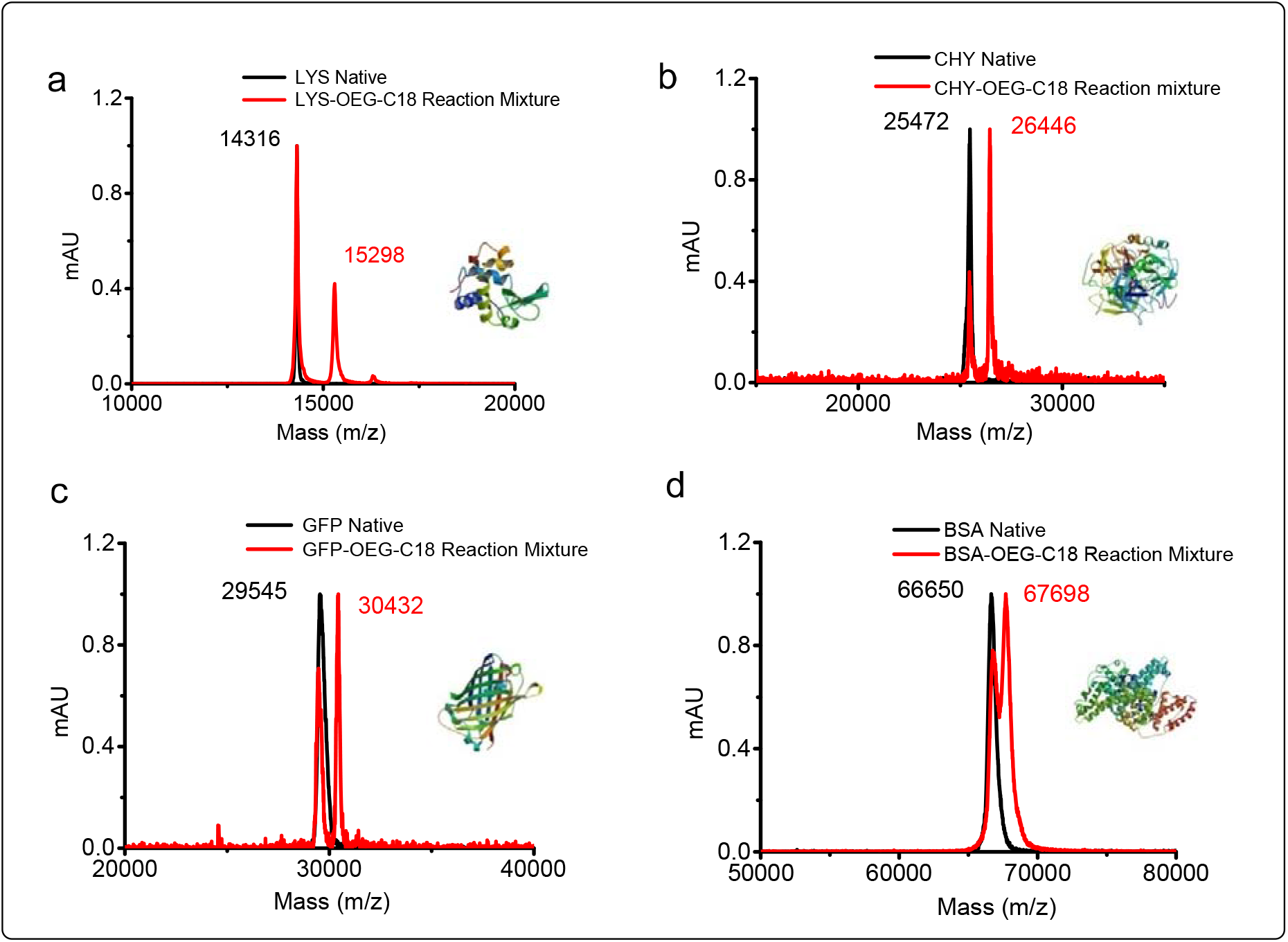
MALDI-ToF MS results **a**. native LYS and a reaction mixture containing the starting material (native LYS) and the product LYS-OEG-C18 **b**. native CHY/reaction mixture **c**. native GFP/reaction mixture **d**. native BSA/reaction mixture. In each case, the successful bioconjugation is evident from appearance of single peak at a higher molecular weight corresponding to the molecular weight of the particular protein bioconjugate.

Although we were successful in a site-specification bioconjugation of a highly hydrophobic synthetic molecule to a globular protein, the next challenge is the purification and analytical characterization of newly designed protein conjugate. These proteins are extremely challenging to purify because of their amphiphilic nature and hence prone to non-specific aggregation. Previously, we had reported an easy method for purification of a diverse set of self-assembling semi-synthetic proteins.^11^ We have adopted the same method for purification. In brief, the Triton-X-100 was removed by ion-exchange chromatography followed by the separation of protein conjugate from native protein using size-exclusion chromatography (SEC). We were delighted to see a single peak for all the bioconjugates manifesting their monodisperse character in mass spec characterization (Figures S2 and S4).

Next, we set out to test whether these bioconjugates can self-assemble into monodisperse protein assemblies. Accordingly, we carried out analytical SEC studies (Figure S7 and S8). We were pleased to see all the protein bioconjugates self-assemble into protein complexes of bigger size as evident from SEC results (Figure 3a-3d). For example, BSA-OEG-C18 oligomerizes to form a 700 kDa protein complex whereas CHY-OEG-C18 and GFP-OEG-C18 formed 400 and 330 kDa protein complexes, respectively. Interestingly, the smallest protein bioconjugate LYS-OEG-C18 also self-assembles to form a trimer of 50 kDa protein nanoreactor (Figure 3a-3d, Table 1). These results are consistent with the previous results from our group that the oligomerization of globular proteins heavily depends on surface charge provided the linker length and hydrophobic tail remain constant.^11a^ This is indeed true in this study as well; LYS, (M_w_ = 14.3 kDa, D_h_ = 2 nm, and pI = 11.35) one of the smallest highly cationic globular protein forms only trimer. However BSA (Mw = 66.6 kDa, D_h_ = 4 nm, pI = 4.7) self-assemble into 700 kDa decamer (Figure 3a-3d, Table 1). Excited by these results, we wanted to check the hydrodynamic size (D_h_) of these protein complexes. Accordingly, we carried out dynamic light scattering studies for BSA-OEG-C18 and CHY-OEG-18 bioconjugates. The hydrodynamic sizes of BSA-OEG-C18 and CHY-OEG-18 are 13 and 12 nm, respectively and these results match with SEC results of this study (Figure 3e, Table 1). Besides, SEC profiles and DLS results of these bioconjugates closely matches with the results obtained for other protein amphiphiles made using active-site bioconjugation technique.^11a^

**Figure 3.**
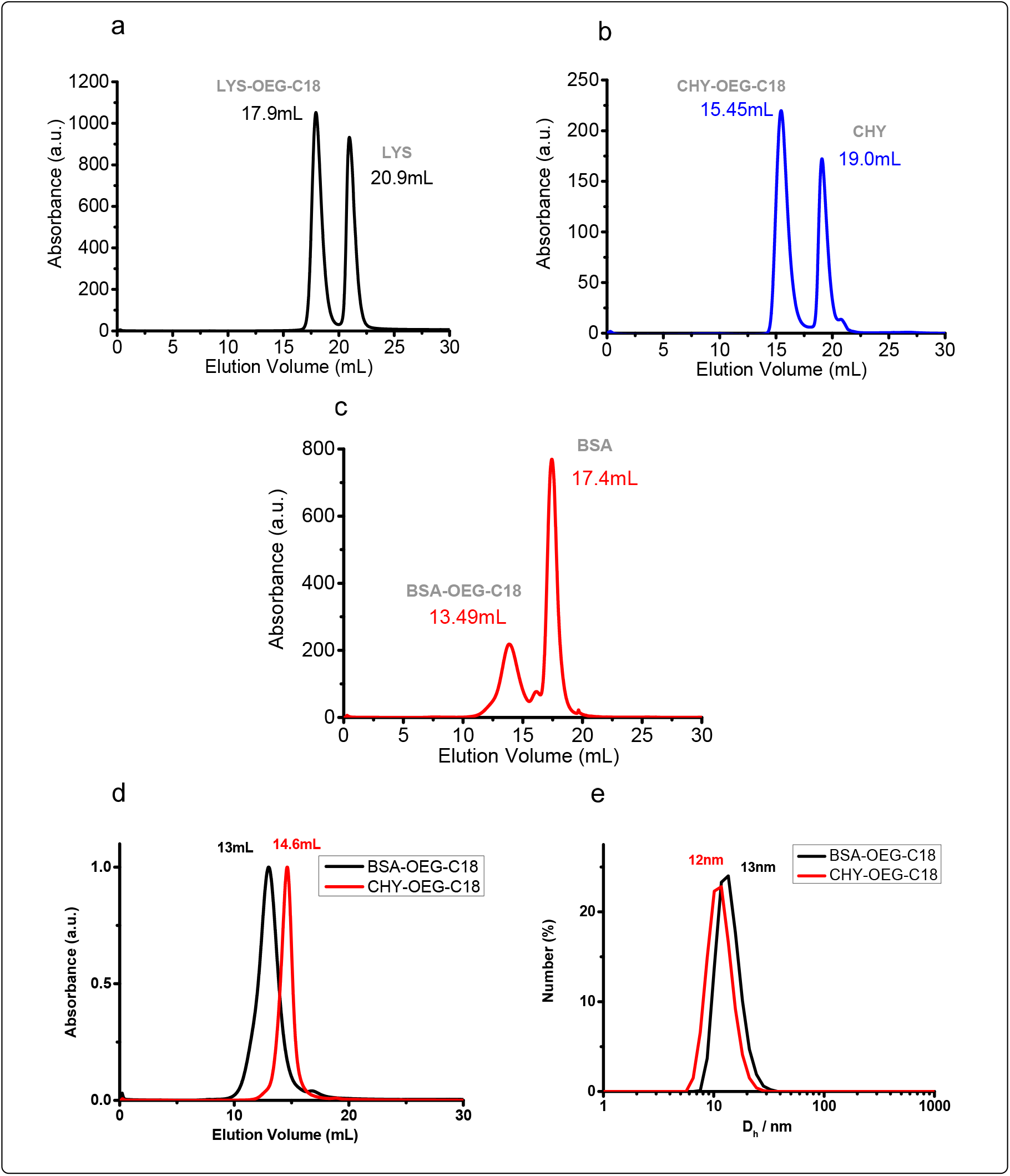
a. SEC chromatogram of bioconjugation reaction mixtures; **a**. LYS-OEG-C18 **b**. CHY-OEG-C18 **c**. BSA-OEG-C18. Semi-synthetic protein bioconjugates self-assemble into larger protein complex and therefore elutes first whereas native protein which exist as a monomeric proteins elutes later **d**. SEC data of purified CHY-OEG-C18 and BSA-OEG-C18 **e**. DLS data of purified CHY-OEG-C18 and BSA-OEG-C18

**Table 1.**
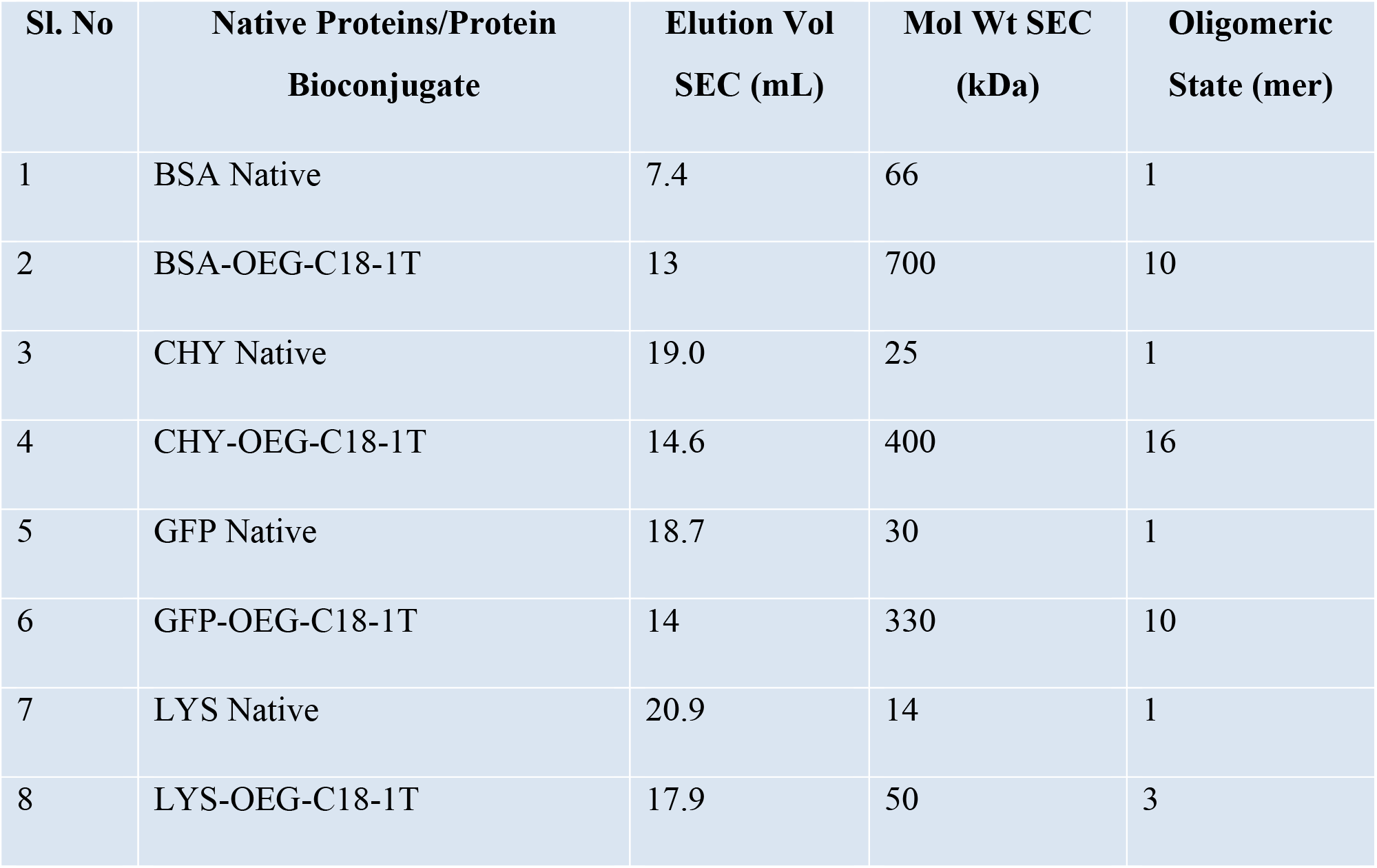
Oligomeric state of native and self-assembling semi-synthetic protein bioconjugates. Molecular weights of native proteins and their corresponding self-assembling semi-synthetic protein complexes were calculated using standard calibration curve (Figure S7 and S8).

The most important objective of this study is to design a protein-based nanoreactor that can mimic the function of a naturally occurring proteasome. It is possible that during self-assembly the active-site of each subunit of CHY-OEG-C18 protein complexes can be sterically hindered and hence diminishes the catalytic activity of the protein by reducing the accessibility of the active-site by the substrate (Figure 4). To check that possibility, we carried out the enzymatic assay of native CHY and CHY-OEG-C18 in parallel. Note that native CHY exists as a monomer whereas CHY-OEG-C18 exists as a decamer (from both SEC and DLS results). For the enzymatic assay, the concentrations of both CHY and CHY-OEG-C18 were kept at 80μM. We chose this concentration because dynamic light scattering studies reveal, CHY-OEG-C18 exists as a 12 nm proteasome-like nanoreactor at 80 μM. The enzyme kinetic results suggest that the rate of hydrolysis of SPNA substrate is comparable between the native CHY and the CHY-OEG-C18 (Figure 4b, 4c). This is a very interesting result considering the catalytic activity of native CHY is not compromised when it oligomerizes into a decamer (Figure 4a). In a way, this process mimics the self-assembly of protease into proteasome which contains 14 subunits of proteases.^2^

**Figure 4.**
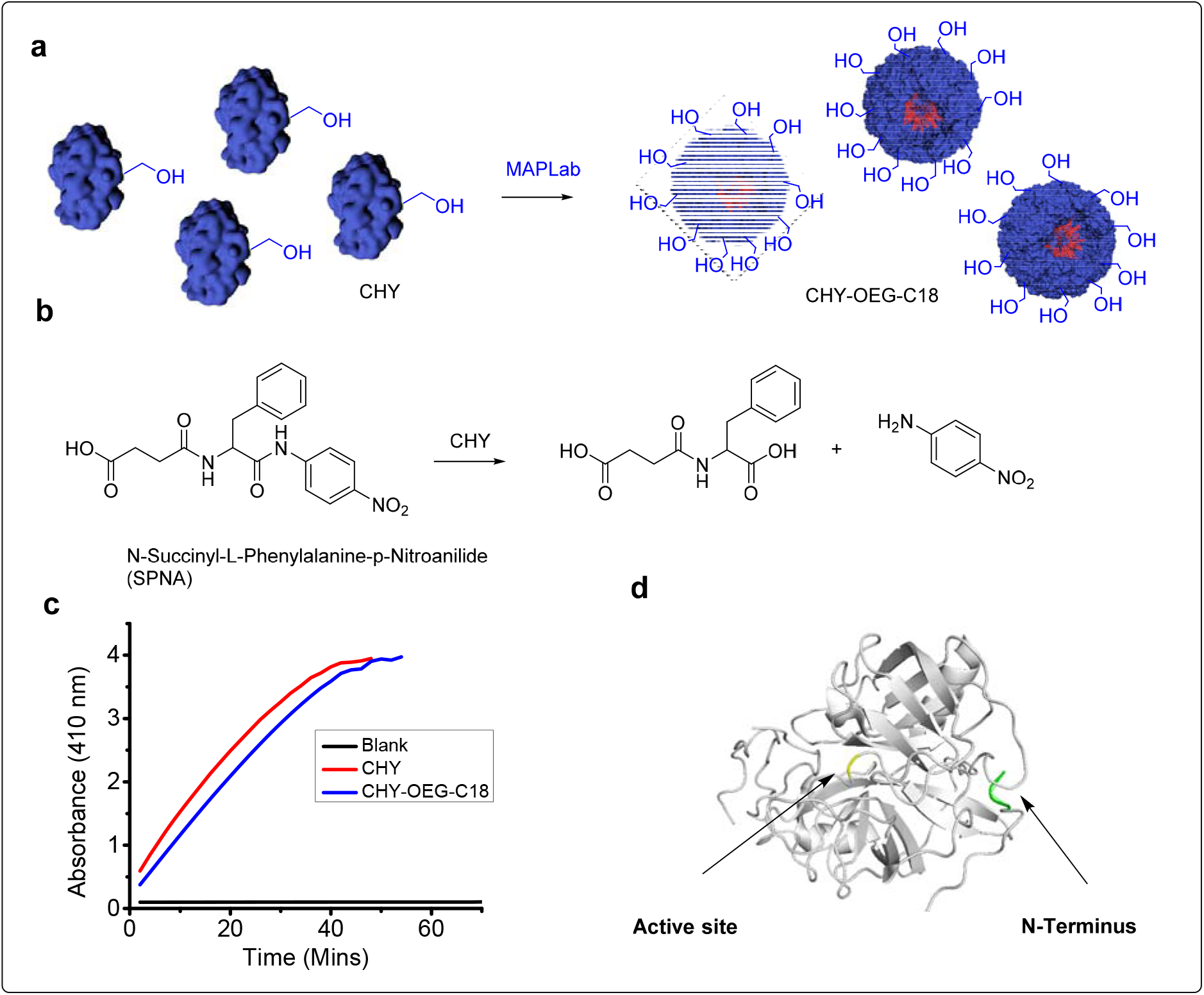
Functional assay. **a**. Conversion of monomeric chymotrypsin (CHY) into proteasome-like nanoreactor using a MAPLab technology. The active site serine residue of chymotrypsin is highlighted in the figure. Conversion of CHY to CHY-OEG-C18 leads to programmed oligomerization to decamer. **b**. Schematic representation of hydrolysis of SPNA substrate by CHY. **c**. Kinetic assay CHY *Vs* CHY-OEG-C18. The rate hydrolysis of SPNA substrate is found to be similar for both monomeric CHY and decamer CHY-OEG-C18. Concentrations of CHY and CHY-OEG-C18 were kept at 80μM.**d**. CHY (PDB accession: 1YPH) structure showing active site serine residue (yellow) with respect to *N*-terminus residue (green).

In summary, we have demonstrated that monomeric CHY can be programmed to oligomerize into monodisperse decamer without compromising on the protease activity whereas LYS can be assembled into a trimer. We have shown here that (i) N-terminal bioconjugation strategy along with MAPLab technology could be used for the design of programmable protein-based nanoreactor; (ii) Both *N*-terminal bioconjugation and self-assembly of protein bioconjugate does not affect the activity of the protein; (iii) the utility of this method with a diverse set of proteins and finally; (iv) the size and oligomeric state of protein complexes can be precisely tuned by choosing an appropriate linker and a hydrophobic tail. The success of this approach depends on the power of two previous technologies i.e. the MAPLab technology^11^ and the *N-*terminal bioconjugation method.^13^ Considering, this is a platform technology, one can envisage using any protease of interest to build semi-synthetic proteasomes. Although the designed protein-based nanoreactor is considered as an amateur compared to naturally occurring proteasomes. This study lays the foundation for the design of semi-synthetic protein-based nanomachines with increased complexity and sophistication. This kind of study also would shed some light on our understanding of the oligomerization of natural proteins. The ability to design the protein-based nanoreactors using a general chemical method opens up tremendous opportunities for the use of these nanoreactors in different applications such as biocatalysis, targeted drug delivery, and *in vivo* imaging.

## Acknowledgments

This work was supported by DST‐SEB Early Career Award to B.S.S. (ECR/2015/000253) and DBT grant (BT/PRI1450/BRB/I0/1370/2015)

