## Supplementary material for "A Universal Chemical Method for Rational Design of Protein-based Nanoreactors": Supp Info

### Experimental section and methods

#### Selection of proteins used in this work

We chose 4 proteins for our study, *i.e.*, LYS (14.3 kDa), CHY (25.4 kDa), GFP (29.6 kDa), and BSA (66.6 kDa). Proteins were purchased from commercial vendors (Except GFP whose His tagged-cloned gene in pET15b vector was generously donated by Dr. Thomas Pucadyil lab), and molecular weights were determined before modification to track the molecular weight changes accurately.

#### Matrix preparation and molecular weight determination

Apart from the molecular weight determination of native proteins, all stages of protein modification and purification were also followed by MALDI-TOF MS. The samples were analyzed in Linear High Mass mode in AB Sciex 4800 plus MALDI-TOF/TOF analyzer with 4000 Series Explorer as software. Mass was scanned between 10,000 Da and 90,000 Da with a focus mass, selected depending on the protein analyzed.

We followed a different MALDI-TOF Ms Procedure in this study. To brief, 15mg of sinapinic acid was weighed in a microcentrifuge tube then add 1.0 ml of matrix solution (70:30 water/acetonitrile with 0.1 % TFA final concentration) was added and vortexed to get the matrix mixture. The sample and matrix mixture were mixed at a 1:10 ratio. When the crystallization observed in centrifuge tubes, 1-2µL of the samples were spotted on the plate and air-dried for 15 minutes. Then the plate was loaded and fired to get accurate molecular weights.

#### Protein modification, monitoring, and purification

Protein modification, monitoring, and purification were done using a similar protocol reported by us previously. However, some changes were made in the equivalents we used and triton X-100 concentrations we used. The AABP was used varied between 25 - 100 eq, which was just 2 eq in our previous studies. Triton X-100, which was used concentrations 100 times (20 mM) more than CMC or 2% of the total volume of the reaction mixture to solubilize the AABPs in the previous study, gave noisy MALDI-TOF Spectra. So we have decreased the Triton X-100 concentration to 10 times more than CMC. Heating the reaction mixture at 37°C triggered precipitation of protein conjugates in the reaction mixtures. So even though we see increased conversions with heating, we did the conjugation at RT.

All the conjugates were purified by two-step purification, *i.e.,* IEX and SEC, performed using FPLC. IEX was performed to remove triton X-100 using either SP sepharose or Q sepharose resins (GE) depending on isoelectric point (pI) and surface charges of proteins. For example, to purify the reaction mixture of LYS (pI: 11.35) and CHY (pI: 8.75) we used SP sepharose, a cation-exchange resin at pH 7.4 and to purify BSA (pI: 4.7) and GFP is (pI: 5.80) we used Q sepharose, an anion-exchange resin at pH 7.4. The obtained IEX fractions were subjected to SEC immediately to remove the native proteins from the protein conjugates in 50 mM sodium phosphate pH 7.4, 1 M NaCl using either Sephacryl S-100 HR 16/60 or Sephacryl S-200 HR 16/60 or Sephacryl S-300 HR 16/60. After SEC the samples were stored at -80 °C

### Synthesis and purification of N-terminus BSA conjugate

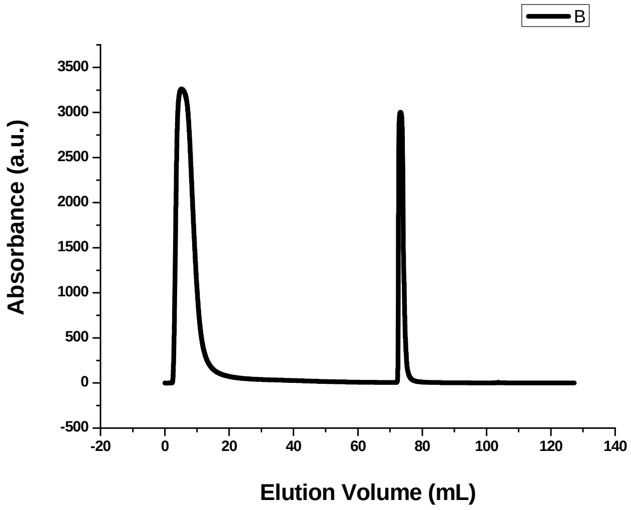


**Figure S1.** IEX chromatogram of BSA reaction mixture.

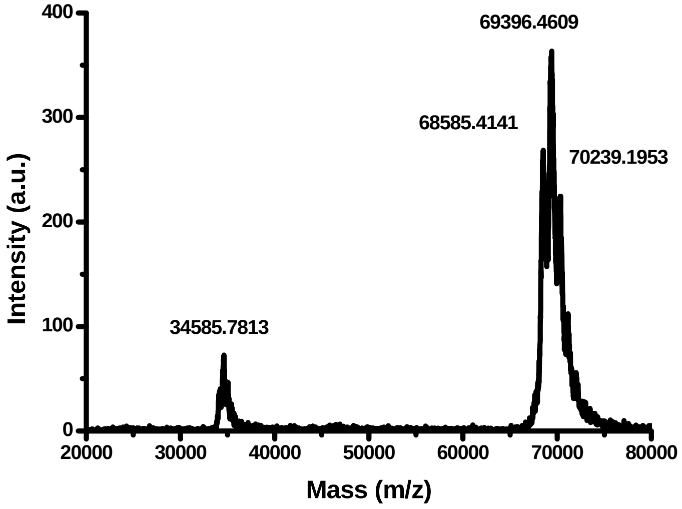


**Figure S2.** MALDI-Tof Spectrum of the purified BSA-OEG-C18.

**Synthesis and purification of N-terminus CHY conjugate**

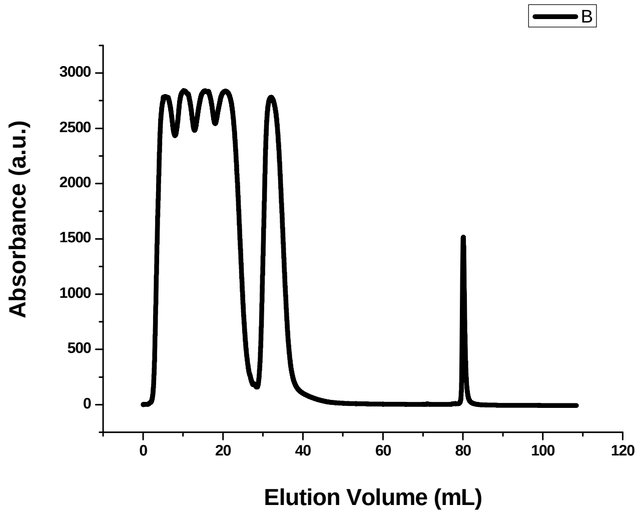


**Figure S3.** IEX chromatogram of the CHY reaction mixture.

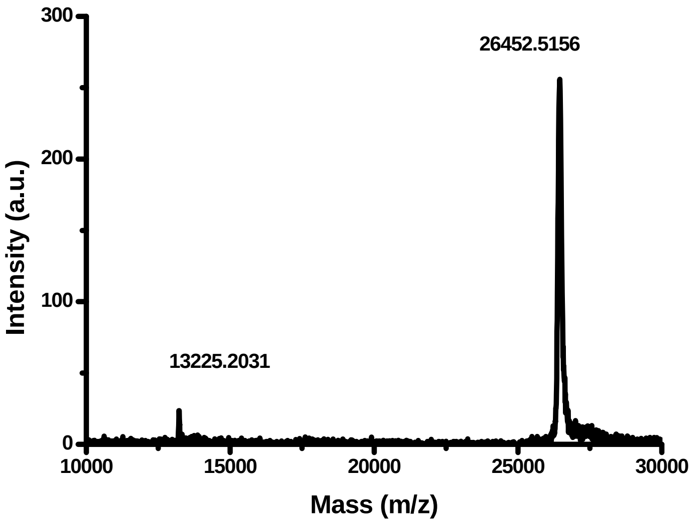


**Figure S4.** MALDI-Tof Spectrum of the purified CHY-OEG-C18.

### Synthesis and purification of N-terminus LYS conjugate

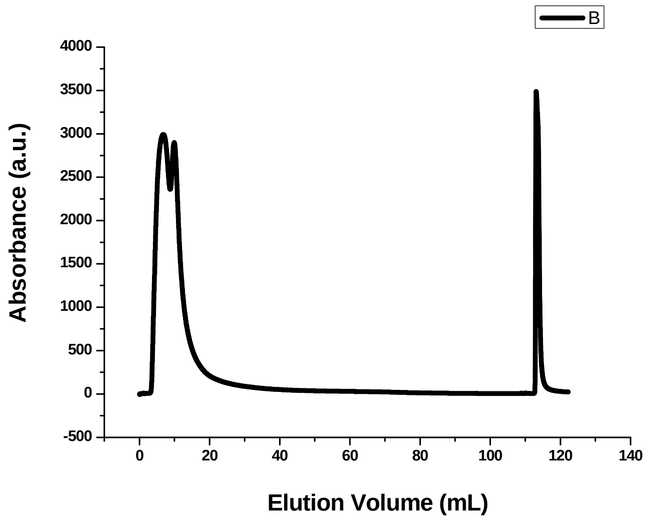


**Figure S5.** IEX chromatogram of LYS reaction mixture.

### Synthesis and purification of N-terminus GFP conjugate

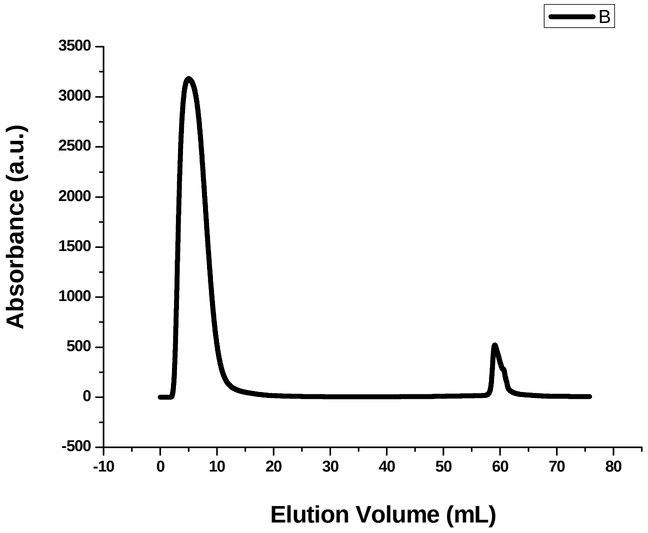


**Figure S6.** IEX chromatogram of GFP reaction mixture.

**Size exclusion-based molecular weight determination**

To determine the molecular weights of soluble protein complexes, we performed SEC in Superose 6 (GE healthcare). For this purpose, we chose standard proteins (GE Healthcare), *i.e.*, blue dextran (2000 kDa), to determine the void volume of this column, thyroglobulin (660 kDa), ferritin (440 kDa), aldolase (158 kDa), conalbumin (75 kDa), ovalbumin (44 kDa), carbonic anhydrase (29 kDa) and trypsin (23 kDa).

A series of size exclusion runs were performed with these proteins in 50 mM sodium phosphate pH 7.4, 200 mM NaCl with 0.25 mL/min as the flow rate. All the proteins were dissolved individually in Milli Q water at 3 mg/mL and 0.5 mL was injected. With the obtained elution volumes, the partition coefficients ($K_{av}$) for standard proteins were calculated according to standard protocol (GE Healthcare) using the formula $K_{av} = V_e - V_o / V_c - V_o$ where, $V_o$ = column void volume (8.5 mL, from the elution volume of blue dextran), $V_e$ = elution volume of the particular sample, and $V_c$ = geometric CV (23.5 mL). The calibration curve was plotted for $K_{av}$ of standards against relative molecular weights (RMW) of standard proteins. ($r^2 = 0.99$, please refer to the calibration curve). Now, to determine the molecular weights of soluble protein complexes, the protein conjugates were dissolved individually in Milli Q water at 5 mg/mL and 0.5 mL was injected.

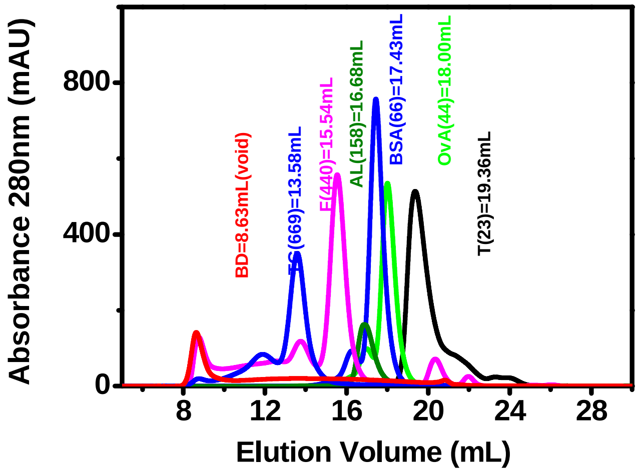


**Figure S7.** SEC runs for standard proteins in Superose 6.

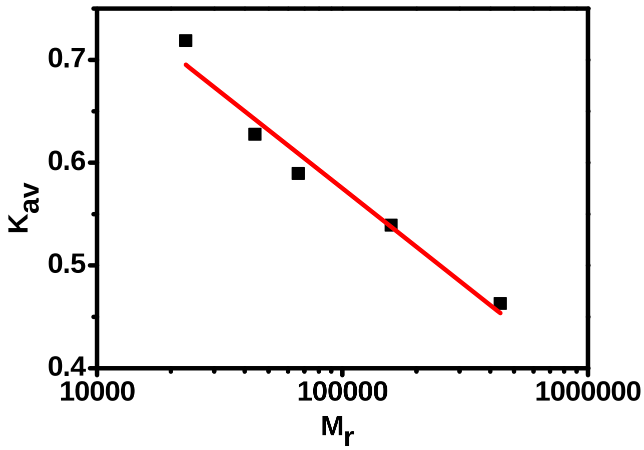


**Figure S8.** Calibration curve for Superose 6.

**Dynamic Light Scattering Studies**

Measurements were carried out at 5 mg/mL concentration in 50 mM sodium phosphate pH 7.4. 1 mL of sample was taken in disposable polystyrene cells and the mean size of the complexes was measured at 90º scattering angle

**Enzymatic Assay**

Enzymatic activity assays were preformed using a microplate reader. All experiments were performed in 5 mM sodium phosphate buffer at pH ) 7.4 with [CHY] ) 80 µM and [CHY-OEG-C18] 80 µM . The enzymatic hydrolysis reaction was initiated by adding a substrate (SPNA) stock solution (16 µL) in DMSO:EtOH (1:9) to a CHY or CHY-OEG-C18 solution (184 µL) to reach a final substrate concentration of 4 mM. Hydrolysis of substrates was monitored for 0-60 min at 405 nm. The assays were performed in triplicate, and the averages are reported.

**Over-expression and purification of GFP-His-tagged protein**

GFP-His tag cloned gene in pET15b vector from Dr. Thomas Pucadiyl Lab was transformed in BL21 (DE3) strain of *E.coli* cells. For transformation, 1 µL (100 ng) of plasmid was added in 25 µL of competent BL21 (DE3) cells and kept at ice for 30 minutes. After incubating it on ice for 30 minutes, heat shock was given for 90 seconds at 42°C in water bath. 950 µL of LB was added and incubated at 37°C for 1 h at thermo mixer, 400 rpm. The transformed cells were centrifuged at 4500 rpm for 5 mins. The pellet obtained after centrifugation was resuspended in 100 µL LB and added on the LB agar plate containing amplicillin antibiotic. The LB agar plated with transformed cells, was then kept at 37°C, incubated for 12 h. After 12 h a single colony was picked and added to 5 mL culture for over-expression.

The protein was expressed in BL21 (DE3) strain of *E.coli* cells (1 L culture) at 37°C to an O.D of 0.6 prior to induction with 0.2 mM IPTG for 3 hours at 18ºC. The cells were pelleted at 6000 rpm for 10 min at 4°C and resuspended in lysis buffer containing 100 mM Tris-HCl, pH-8, 100 mM NaCl, 20 mM imidazole and lysed using sonicator. After sonication, the sample was centrifuged for 30 min at 20,000 rpm.

After removal of cellular debris the supernatant was incubated with Ni-NTA column pre-equilibrated with buffer A (100 mM Tris-HCl, 100 mM NaCl and 20 mM imidazole). The column was washed with 8 bed volume of buffer A twice. After incubating the column with buffer A, protein lysate was loaded in Ni-NTA column. Column incubated with protein lysate was then washed with 10 column volume of buffer B (10 mM Tris-HCl, 500 mM NaCl and 20 mM imidazole) twice. Protein was then eluted within 1 mL fractions with 6 mL of elution buffer, Buffer C (50 mM Tris-HCl, pH-8, and 150 mM NaCl and 100 mM imidazole). Fractions were checked on SDS-PAGE for purity. This was followed by dialysis against buffer A. The protein was concentrated and stored at -80°C in (10 mM Tris-HCl, 300 mM NaCl, 10% glycerol).

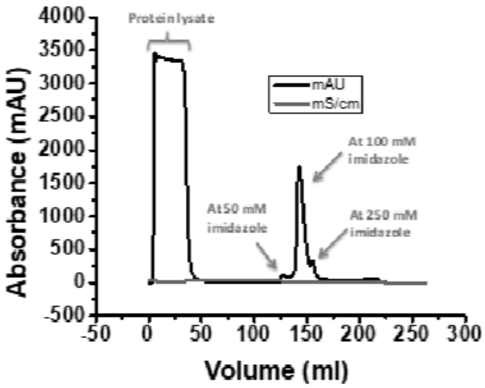


**Fig.S9.** FPLC Chromatogram of the protein lysate loaded onto the Ni-NTA column. The broad peak (0-50 mL elusion volume) is the unbound protein eluted at 20 mM imidazole concentration and sharp peak (~ 150 mL elusion volume) is the GFP-His tag protein bound to the Ni-NTA column eluted at 100 mM imidazole concentration.

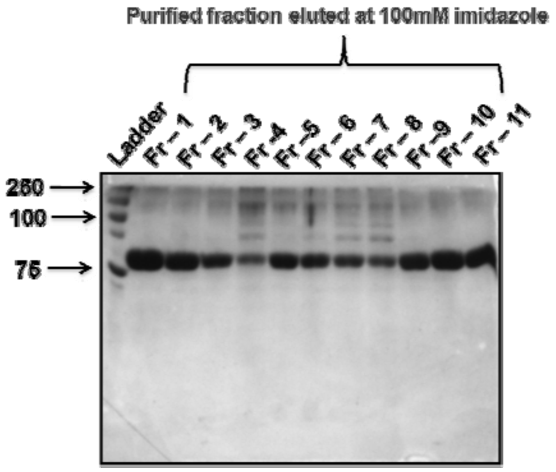


**Fig S10.** SDS-PAGE analysis of Ni-NTA affinity column purified GFP-His tagged protein. Protein samples were run on 12.5% SDS-PAGE gel and stained with coomassie brilliant blue. Lane 1; Ladder; Lane 2-12 are the purified fractions eluted with 100 mM imidazole.

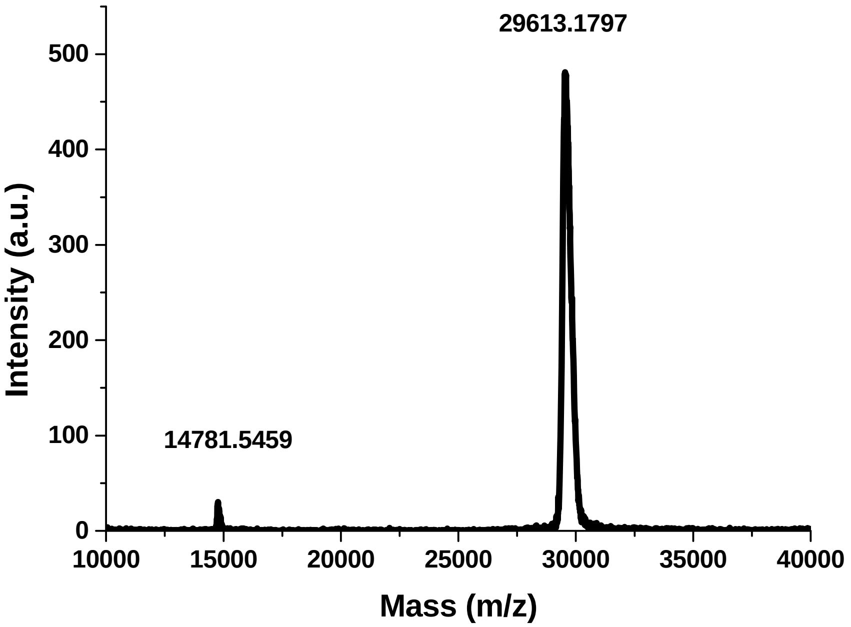


**Fig S11.** GFP- His tagged protein was confirmed with the peak obtained at 29613 Da.

**Synthesis, purification and Characterization of Intermediates and Target Probe**

**Compound 2**

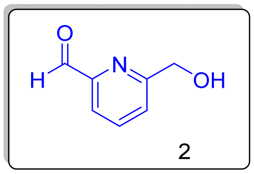

Mol. formula: $C_7H_7NO_2$

Mol. Weight: 137.13 g/mol

Physical appearance: white solid

Yield: 60 %

To a solution of 2,6-pyridinedimethanol (500 mg, 3.62 mmol) in 1,4-dioxane (10 mL), $SeO_2$ (200 mg, 1.81 mmol) was added. The resulting mixture was sonicated for 5 mins and then stirred at 65 □C for 24 hrs. Then the reaction was cooled to room temperature and diluted with dichloromethane. The mixture was filtered through celite, and the filtrate was concentrated under reduced pressure. The resulting crude material was purified by chromatography using PE/EtOAc to get a liquid that became off-white solid later (300mg, 60%). **$^1$H NMR** (400 MHz, $CDCl_3$): $\delta_H$ 10.02 (s, 1H), 7.84 (m, 2H), 6.52 (m, 1H), 4.48 (s, 1H). **$^{13}$C NMR (**100 MHz, $CDCl_3$): $\delta_C$ 193.12, 160.43, 151.66, 137.86, 125.00, 120.65, 77.16, 64.24.

**Compound 3**

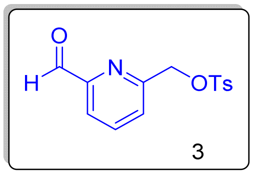

Mol. formula: $C_{14}H_{13}NO_4S$

Mol. weight: 291.32 g/mol

Physical appearance: white solid

Yield: 55 %

Tosyl chloride (542 mg, 2.64 mmol) was added to the solution of alcohol (**compound 42**) (300 mg, 2.2 mmol) in DCM (10mL) at 0 □C. $Et_3N$ (661 mg, 906 µl, 6.54 mmol) was then added to the above mixture, maintaining the temperature at 0 □C. After 1hour at RT, the reaction mixture was concentrated and purified using column chromatography using PE / EtOAc (350mg, 55%). **$^1$H NMR** (400 MHz, $CDCl_3$): $\delta_H$ 9.93 (s, 1H), 7.85 (m, 4H), 7.66 (d, 1H), 7.35 (d, $J$ = 8.4 Hz,

2H), 5.22 (s, 2H), 2.43 (s, 3H). **$^{13}$C NMR (**100 MHz, $CDCl_3$): $\delta_C$ 154.80, 152.24, 145.41, 138.22, 132.67, 130.09, 128.18, 126.10, 121.35, 77.16, 71.19, 21.75. **HRMS** (MW+H): 292.06 g/mol.

**Compound 6**

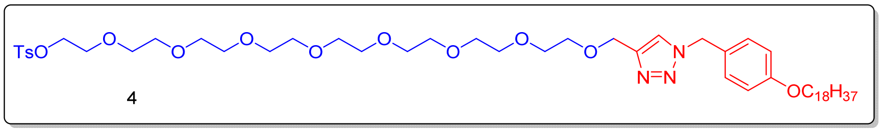


Mol. formula: $C_{51}H_{85}N_3O_{12}S$

Mol. weight: 964.31 g/mol

Physical appearance: yellow waxy solid

Yield: 75 %

Synthesis of alkyne (**compound 4**) and azide (**compound 5**) was done as per our previous report. Alkyne (5 g, 8.89 mmol) and azide (2.82 g, 8.89 mmol) were weighed in an oven dried RBF and THF was added, followed by water with vigorous stirring. Then, Na Asc. (6.6 mg, 0.033 mmol) was added followed by $CUSO_4$ (2.6 mg, 0.016 mmol) and allowed to react overnight. The reaction was extracted with DCM and then concentrated under reduced pressure. The resulting crude material was purified by NPC using PE/EtOAc (6.4 g, 75 %). **$^1$H NMR** (400 MHz, $CDCl_3$): $\delta_H$ 7.78 (d, $J$ = 8.4, 2H),7.47 (s, 1H), 7.33 (d, $J$ = 8, 2H), 7.20 (d, $J$ = 6.8 Hz, 2H), 6.86 (d, $J$ = 8 Hz, 2H), 5.41 (s, 2H), 4.62 (s, 2H), 4.1 (m, $J$ = 6.8Hz, 2H), 3.92 (t, $J$ = 6.4 Hz, 2H), 3.63 (m, 30H), 3.5 (m, 5H), 1.75 (t, $J$ = 7.2 Hz, 2H), 1.42 (m, 2H), 1.24 (m, 34H), 0.86 (t, $J$ = 6.4 Hz, 3H). **$^{13}$C NMR (**100 MHz, $CDCl_3$): $\delta_C$ 159.57, 144.89, 133.11, 129.79, 128.07, 126.37, 115.05, 77.16, 70.62, 69.78, 69.49, 68.75, 68.19, 32.01, 29.78, 29.69, 29.67, 29.48, 29.45, 29.28, 26.11, 22.78, 21.74, 14.23. **MALDI-TOF MS** (MW+ Na): 986.55 g/mol.

**Compound 7**

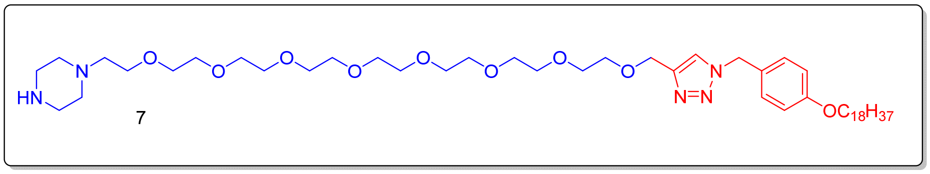


Mol. formula: $C_{48}H_{87}N_5O_9$

Mol. weight: 878.25 g/mol

Physical appearance: yellow waxy solid

Yield: 82 %

Tosylate (**compound 6**) (2 g, 2.07 mmol) and piperzine (1.9 g, 22.09 mmol) were dissolved in THF (30 mL). Then the mixture was refluxed at 65 □C. After 12 hrs, reaction mixture was concentrated and purified by NPC (1.5 g, 82 %). $^{1}$**H NMR** (400 MHz, $CDCl_3$): $\delta_H$ 7.48 (s, 1H), 7.12 (d, $J$ = 8.8 Hz, 2H), 6.77 (d, $J$ = 8.8 Hz, 2H), 5.43 (s, 2H), 4.53 (s, 2H), 3.82 (t, $J$ = 6.4 Hz, 2H), 3.51 (m, 31H), 3.04 (t, $J$ = 4.4 Hz, 4H), 2.51 (m, 5H), 1.66 (m, 2H), 1.34 (m, 3H), 1.15 (m, 31H), 0.769 (t, $J$ = 6.8 Hz, 3H). $^{13}$**C NMR (**100 MHz, $CDCl_3$): $\delta_C$ 159.28, 145.12, 129.51, 126.20, 122.23, 114.77, 77.16, 70.36, 70.17, 69.50, 68.58, 68.34, 67.90, 64.47, 57.48, 57.17, 53.43, 50.12, 45.72, 43.55. **MALDI-TOF MS** (MW+ Na): 900.72 g/mol.

**Compound 8**

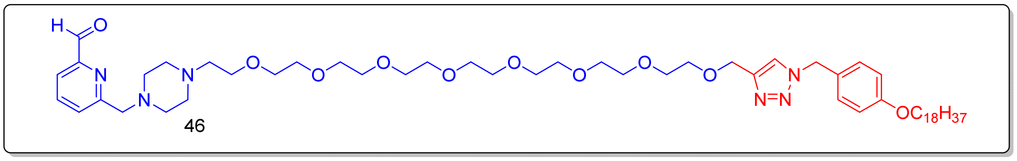


Mol. formula: $C_{55}H_{92}N_6O_{10}$

Mol. weight: 997.37 g/mol

Physical appearance: yellow waxy solid

Yield: 33 %

**Compound 7** (1.6 g, 1.82 mmol), $K_2CO_3$ (0.500 g, 3.62 mmol) and tosylate (**compound 3**) (0.530g, 1.82 mmol) were weighed in RBF. Then the mixture was dissolved in ACN (10 mL) and refluxed at 65 □C. After 16 hrs, reaction mixture was concentrated and purified by NPC (0.6 g, 33 %). $^{1}$**H NMR** (400 MHz, $CDCl_3$): $\delta_H$ 10.06 (s, 1H), 7.83 (m, 2H), 7.8 (m, 1H), 7.43 (s, 1H), 7.21 (d, $J$ = 8.4 Hz, 2H), 7.67 (d, $J$ = 8.4, 2H), 5.42 (s, 2H), 4.63 (s, 2H), 3.92 (t, $J$ = 6.4 Hz, 2H), 3.75 (s, 2H), 3.61 (m, 32H), 2.59 (m, 9H), 1.75 (m, 2H), 1.42 (m, 2H), 1.24 (m, 32H), 0.86 (t, $J$ = 6.8 Hz, 3H). $^{13}$**C NMR (**100 MHz, $CDCl_3$): $\delta_C$ 193.77, 137.51, 129.81, 120.29, 115.08, 77.16, 70.67, 70.48, 68.23, 64.91, 64.17, 57.80, 53.83, 53.68, 53.34, 32.04, 29.81, 29.61, 29.31, 26.14, 22.81, 14.25. **MALDI-TOF MS** (MW+ Na):1019.60 g/mol.
